# MDH2 is a metabolic switch rewiring the fuelling of respiratory chain and TCA cycle

**DOI:** 10.1101/2021.04.17.440272

**Authors:** Thibaut Molinié, Elodie Cougouilles, Claudine David, Edern Cahoreau, Jean-Charles Portais, Arnaud Mourier

## Abstract

The mitochondrial respiratory chain (RC) enables many metabolic processes by regenerating both mitochondrial and cytosolic NAD^+^ and ATP. The oxidation by the RC of the NADH metabolically produced in the cytosol involves redox shuttles as the malate-aspartate shuttle (MAS) and is of paramount importance for cell fate. However, the specific metabolic regulations allowing mitochondrial respiration to prioritize NADH oxidation in response to high NADH/NAD^+^ redox stress have not been elucidated. Our work demonstrates the crucial role played by MDH2 in orchestrating the electron fuelling of the RC. On one hand, MDH2 mediated oxaloacetate (OAA) production allows mitochondria to metabolize pyruvate and glutamate and to feed complex I with NADH, whereas on the other hand, OAA inhibits complex II. This regulatory mechanism synergistically increases RC’s NADH oxidative capacity and rewires MDH2 driven anaplerosis of the TCA favouring the oxidation of malate produced by the MAS.

## INTRODUCTION

Mitochondria are organelles present in almost all eukaryotic cells and are essential for the maintenance of cellular metabolism. The oxidative phosphorylation system (OXPHOS), which is located in the inner mitochondrial membrane (IMM), is composed of two functional entities: the respiratory chain (RC) and the phosphorylation system, which includes the ATP synthase and carriers, such as the ATP/ADP carrier and the phosphate carrier (Chance and Williams, 1956). Historically, the mitochondrial respiratory chain has been defined as an ensemble of four protein complexes (I, II, III, and IV) and of mobile electrons carriers as coenzyme Q (CoQ) and cytochrome-*c* (Cyt *c*). The RC complexes I, III and IV are defined as coupling sites as they couple redox reactions to proton translocation outside the matrix space. The protons electrochemical potential generated across the IMM by the RC, allows the ATP synthase (complex V) to generate ATP (Mitchell, 1961). Beyond ATP synthesis, mitochondrial maintenance of NADH/NAD^+^ redox cofactor balance is of paramount importance for numerous mitochondrial and cytosolic metabolic pathways (Alkan et al., 2020; Borst, 2020; Lee et al., 2020). Redox shuttles, such as the Malate Aspartate Shuttle (MAS), or the *α*-glycerophosphate shuttle are required to allow mitochondrial RC to indirectly oxidize the cytosolic NADH produced by glycolysis and other metabolic pathways (Cederbaum et al., 1973; Dawson, 1979; Safer et al., 1971). The MAS has been demonstrated to be the predominant NADH redox shuttle in highly oxidative tissues as brain, heart and liver (LaNoue et al., 1974, 1973). This key redox shuttle involves mitochondrial carriers as well as cytosolic and mitochondrial malate dehydrogenases (MDH1, MDH2) and glutamate aspartate transaminases (GOT1, GOT2) (Borst, 2020). Intercompartment metabolite cycling through the MAS allows MDH1 and MDH2 to work in opposite directions (oxidizing cytosolic NADH and reducing mitochondrial NAD^+^) enabling complex I to indirectly oxidize cytosolic NADH. Beyond NADH redox maintenance through Complex I, the RC is also key to sustain the tri-carboxylic acid cycle (TCA cycle) via complex II. This enzyme has recently emerged as focal point of investigation in neurodegeneration (Baysal et al., 2000; Browne et al., 1997), cardiovascular disorders (Chouchani et al., 2014; Dröse, 2013; Zhang et al., 2018), immunology (Mills et al., 2016), cancer biology (Favier et al., 2015; King et al., 2006; Selak et al., 2005) and signalling (Finley et al., 2011; He et al., 2004; Zhang et al., 2011).

The fact that both complex I and II rely on CoQ as common substrate, prompted scientists to investigate molecular processes allowing the RC to orchestrate the activity of these two CoQ-oxidoreductases. A recent burst of interest in understanding regulation and activity of CoQ oxidoreductases was triggered by evidence that in contrast to Complex II, the vast majority of Complex I physically interact with other RC complexes III and IV (Schägger and Pfeiffer, 2000). One of the most important arguments for a higher order organization of the respiratory chain was provided by the use of blue native polyacrylamide gel electrophoresis (BN-PAGE), which showed that respiratory chain complexes from a wide range of organisms can be extracted in supramolecular assemblies when mitochondria are solubilized using mild detergents (Schägger and Pfeiffer, 2000). In mammalian mitochondria, supercomplexes (SCs) consisting of complexes I, III, and IV are functional entities consuming oxygen in presence of NADH and are therefore named respirasome (Acín-Pérez et al., 2008; Dudkina et al., 2010; Schägger and Pfeiffer, 2000). Recent studies using a multitude of independent approaches, such as electron cryo-microscopy (cryo-EM) (Gu et al., 2016; Letts et al., 2016), electron cryotomography (cryo-ET) (Davies et al., 2018), cross-linking mass spectrometry (Chavez et al., 2018), and Förster resonance energy transfer (FRET) analyses (Rieger et al., 2017), have convincingly proven the existence of respirasomes in various organisms. Respirasomes have been reported to play a critical role in bioenergetics and proposed to sequester dedicated pools of CoQ and Cyt *c* in order to function as independent bioenergetic units. It has also been proposed that respirasome could reduce the diffusion distance of CoQ and Cyt *c*, thereby increasing the electron transfer efficiency between complexes organized in respirasomes (Balsa et al., 2019; Bianchi et al., 2004; Lapuente-Brun et al., 2013). However, analyses of supercomplex structures, as well as of kinetic and spectroscopic data, do not support substrate channelling by sequestration of CoQ and Cyt *c* (Blaza et al., 2014; Fedor and Hirst, 2018; Trouillard et al., 2011). Therefore, the bioenergetic roles of respirasomes on mitochondrial NADH oxidation efficiency and on prioritizing electron delivered by complex I against the respirasome-free complex II remain in question.

A growing body of evidence suggests that intramitochondrial oxaloacetate (OAA) level could play a key role in orchestrating the fuelling of the RC from complex I or II. The OAA was identified decades ago as a potent complex II inhibitor (Oestreicher et al., 1969; Schollmeyer and Klingenberg, 1961; Wojtczak et al., 1969; Zeylemaker et al., 1969). However, whether or not OAA inhibition of complex II is a metabolically and physiologically relevant process remains controversial. The lack of progress on this essential question most probably lies in the numerous technical constraints, precluding the proper characterization of the OAA metabolic and compartmentation complexity, as well as its chemical instability preventing its quantification by mass spectrometry. Interestingly, recent reports from the Sivitz laboratory characterized mitochondria isolated from different mouse tissues and demonstrated that OAA inhibition of complex II was strongly controlled by mitochondrial membrane potential (Fink et al., 2019, 2018).

In this study, we carefully characterize the bioenergetic properties of heart and liver murine mitochondria, as their metabolism strongly depend on MAS (LaNoue et al., 1973; Safer B, 1975), but also differ in their respective capacity to oxidize NADH versus succinate (Brandt et al., 2017). Using metabolically active mitochondria, we developed independent biochemical approaches to decipher the respective role of RC supramolecular organisation as well as OAA driven complex II inhibition in orchestrating the electron fuelling of the RC from complex I and II. Our results demonstrate that despite the kinetic advantage potentially conferred to NADH oxidation by the assembly of complex I into respirasomes, liver and heart RC prioritize electron provided by complex II. Interestingly, our analyses enlighten the crucial role played by MDH2 in orchestrating electron fuelling of the RC. On one hand, MDH2 mediated OAA production is indispensable to allow mitochondria to metabolize pyruvate and glutamate and to feed complex I with NADH, whereas on the other hand, OAA produced by MDH2 inhibits complex II. Remarkably, the MDH2 mediated rewiring RC electrons flow priority from succinate toward NADH oxidation is triggered by physiological concentration of L-malate. Consequently, oxidation of imported L-malate by MDH2 does not only rewire the fuelling of the RC from Complex II toward complex I, but concomitantly rewire anaplerosis of TCA cycle, downregulating malate produced by the fumarase to favour oxidation of imported malate. Consequently, MAS does not just passively balance NADH/NAD^+^ redox between cytosol and mitochondria. Instead, extramitochondrial L-Malate produced by the MAS in response to NADH redox stress can, once imported inside mitochondria, rewires RC electron flow and TCA cycle activity to increase mitochondrial NADH oxidation capacity.

## RESULTS

### Heart and liver mitochondria strongly differ in their NADH and succinate oxidative capacities

To elucidate how mitochondrial RC orchestrates NADH and succinate oxidation we characterized heart and liver mitochondria presenting high TCA and MAS metabolic activities. Bioenergetic characterization of isolated mitochondria was first performed by measuring the oxygen consumption under phosphorylating, non-phosphorylating and uncoupled conditions using high-resolution O2K oxygraphs. To determine the capacity of heart and liver mitochondria to oxidize NADH or succinate, metabolically active mitochondria were incubated in the presence of various nutrient combinations whose metabolism results in delivery of electrons at the level of complex I or complex II. In both heart and liver, we found that simultaneous addition of pyruvate, glutamate and malate was the best combination of nutrient to feed complex I with electrons (Figure S1A-B). Then, we confirmed that adding complex I inhibitor (rotenone) in presence of succinate was essential to assess maximal complex II driven respiration (Figure S1C-D). In line with previous observations (Brandt et al., 2017), respirations assessed under phosphorylating or uncoupled conditions were five to three times higher in heart than in liver mitochondria (Figure 1A-B). Furthermore, our analyses showed that, compared to the succinate driven respiration, heart mitochondria present higher complex I driven respiration (Figure 1A) than liver mitochondria (Figure 1B). This observation prompted us to decipher if the difference in complex I and II driven respirations, was caused by mitochondrial metabolic constraint affecting NADH or succinate provisional of the RC or to intrinsic RC functional discrepancies. To this end, we permeabilized mitochondria to specifically measure RC activity fed directly with NADH or succinate. Interestingly, when RC is directly fuelled with reduced substrates, heart RC can oxidize NADH two time faster than succinate (Figure 1C) whereas an opposite situation is observed in liver RC (Figure 1D). The high disparity in NADH and succinate oxidation rates observed between heart and liver RC prompted us to further characterize tissue specific RC composition changes. To this end, we quantified RC complex levels assessing cytochromes (Figure 1E) and multiple OXPHOS protein levels (Figure 1F-H). These two independent approaches consistently showed that all cytochromes as well as RC complex subunits were two to four times more abundant in heart than in liver mitochondria (Figure 1E-G). Moreover, the relative abundance of complex I normalized to complex II was almost tripled in heart compared to liver (Figure 1H). Altogether, the data presented in Figure 1 show that RC complexes content and stoichiometry strongly differ between heart and liver (Figure 1E-H) and that these differences correlate with tissue specific functional differentiation (Figure 1A-D). Interestingly, when both complex I and complex II substrates are added to mitochondria, the increase in RC activity is too low to sustain both NADH and succinate oxidation flow at maximal speed (Figure 1A-D). This observation prompted us to develop assays allowing to determine the respective contribution of complex I and complex II in feeding RC with electrons, when both NADH and succinate are present.

**Figure 1:**
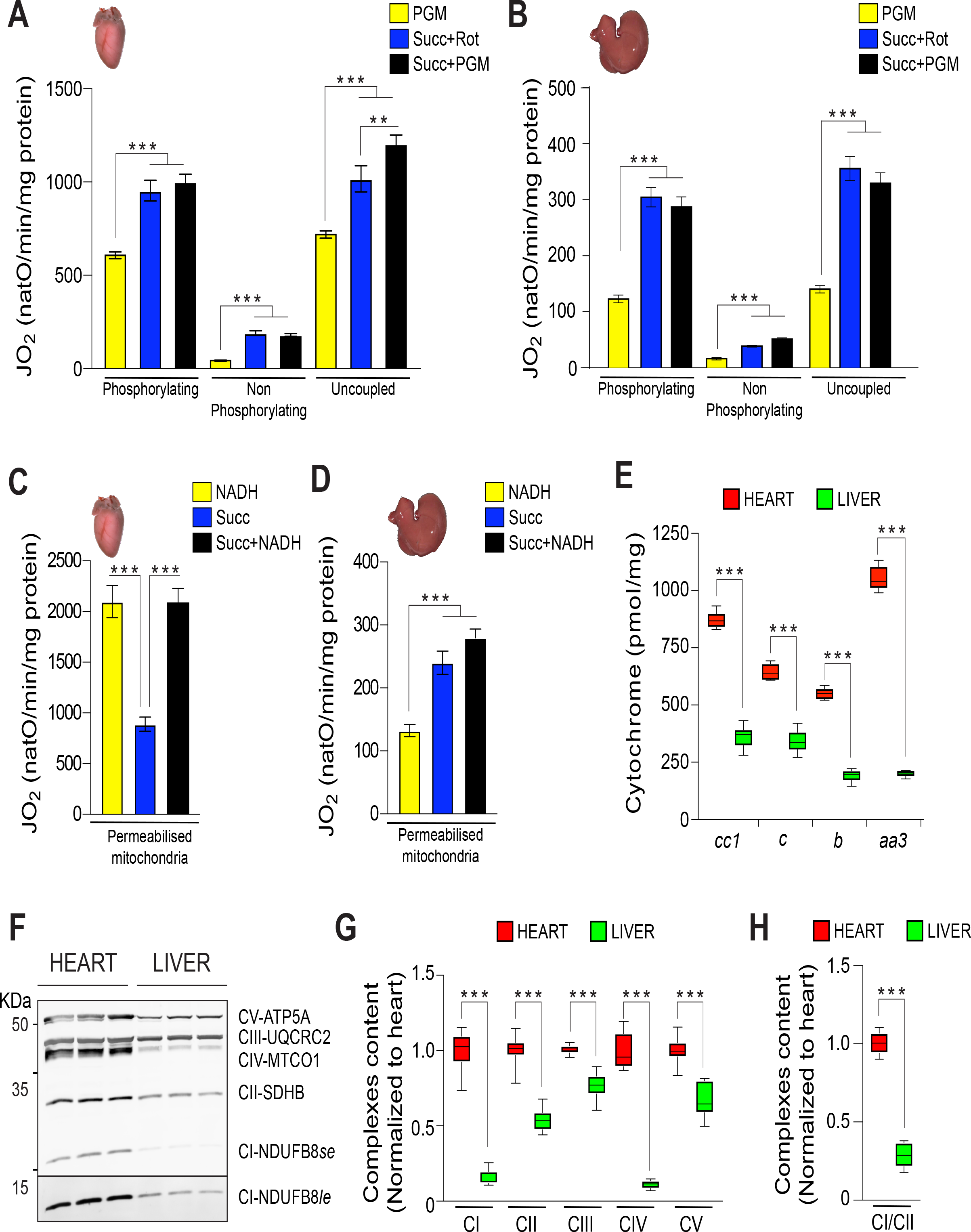
Heart and liver respiratory chain composition and bioenergetics. (A, B) Oxygen consumption of intact heart (A) and liver (B) mitochondria assessed in presence of mitochondrial complex I substrates (PGM : pyruvate 10 mM, glutamate 5 mM, malate 5 mM), or complex II substrate (Succ+Rot : Succinate 10 mM and rotenone 30 nM), and PGM and succinate (Succ+PGM). Respiration are assessed under phosphorylating conditions (ADP+Pi), non-phosphorylating conditions (oligomycin), and uncoupled (CCCP titration). Heart (n= 15), liver (n= 5), error bars represent the mean ± SEM. (C, D) Oxygen consumption of permeabilised heart (C) and liver (D) mitochondria assessed in presence of NADH (1.2 mM), Succinate (10 mM) and cytochrome *c* (62.5 µg/ml). Heart (n= 10), liver (n= 12), Error bars represent the mean ± SEM (E) Mitochondrial cytochrome quantified from redox absorbance spectra. Cytochromes content of RC complexes: complex III (*b*, *c1*), complex IV (*aa3*) and cytochrome *c* (*c*) were determined in mitochondrial extracts of heart and liver. Heart (n= 6) liver (n= 6), error bars represent the mean ± SEM. (F) Steady-state levels of OXPHOS subunits, in heart and liver mitochondria, determined by western blot analyses. Short and long exposure are performed to detect complex I (complex I *se* and complex I *le*) in the liver. Representative Western-blot of five different experiments. (G) Relative amount of OXPHOS complexes (complex I– complex V) in heart and liver mitochondria determine by densitometric quantification of western blot experiments presented in (F), and normalized to heart OXPHOS complexes. (n= 5) Error bars represent the mean ± SEM (H) Relative amount of complex I normalized to complex II in heart and liver mitochondria determined by densitometric analyses, (n= 5) Error bars represent the mean ± SEM.

### Heart and liver respiratory chain preferentially oxidize succinate over NADH

The respective contribution of NADH and succinate oxidation in fuelling RC with electrons when mitochondria are in presence of both substrates, was first evaluated using validated and highly specific complex I or II inhibitors *i.e.* rotenone (Rot) and Atpenin A5 (AtpnA5) (Figure S2A-D). Interestingly, this inhibitor driven approach consistently demonstrates that when permeabilized heart and liver mitochondria are incubated with both NADH and succinate, complex II driven respiration was preserved whereas complex I driven respiration was strongly and significantly affected (Figure 2A-B). This significant downregulation of complex I activity in presence of succinate, was confirmed by enzymatic measurement of the NAD^+^ production rate (Figure 2C-D). Interestingly, despite the great disparity in their respective complex I and II oxidative capacities (Figure 1C-D), the priority given to complex II electrons, was consistently observed in both heart and liver mitochondria (Figure 2A-D).

**Figure 2:**
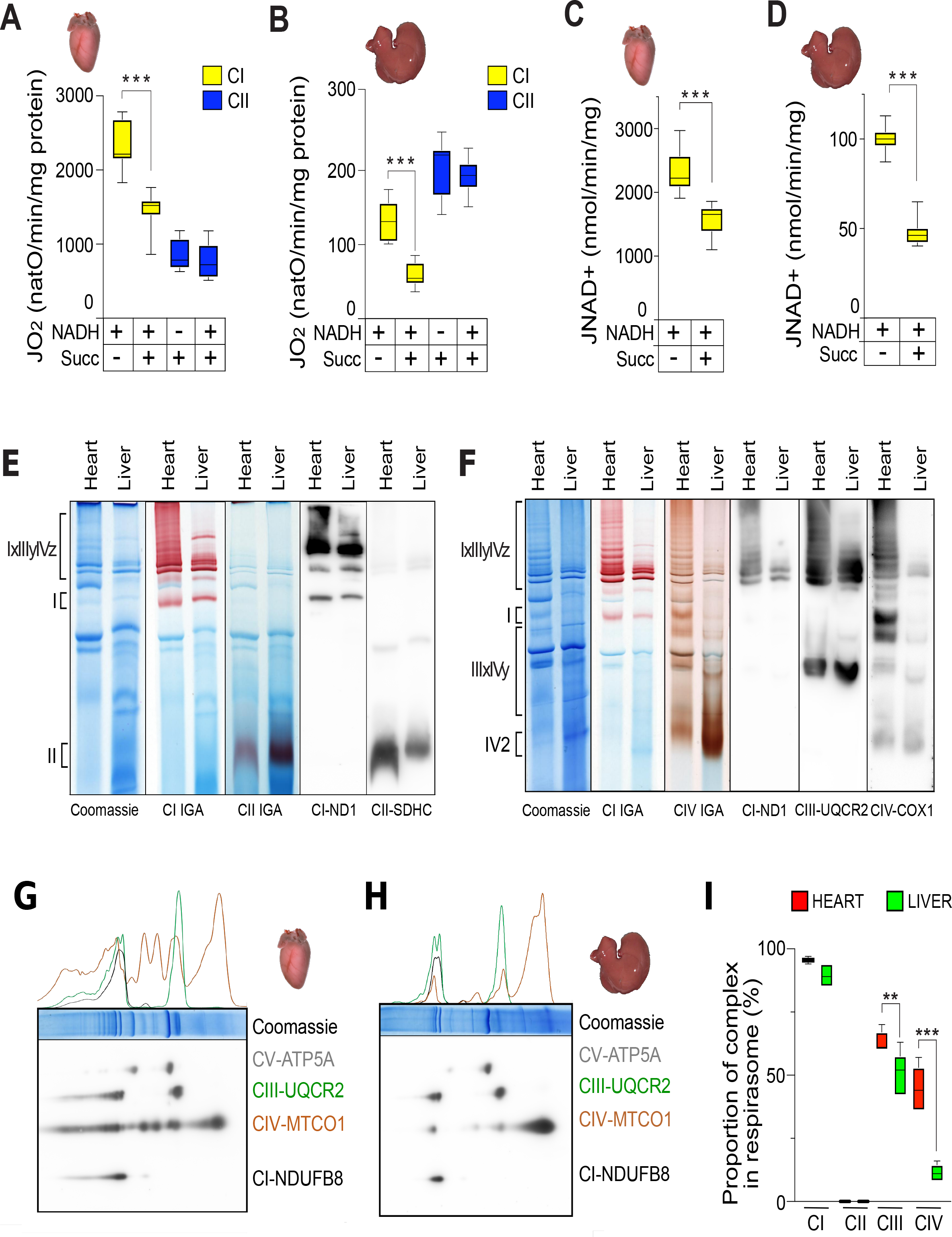
Heart and liver respiratory chain supramolecular organization and preferential electron fuelling. (A, B) Oxygen consumption of permeabilised mitochondria isolated from heart (A) and liver (B) assessed in presence of cytochrome *c* (62.5 µg/ml), complex I substrate NADH (1.2 mM), complex II substrate succinate (Succ, 10 mM) or both substrates simultaneously added. Addition of complex I (rotenone, 30 nM) or complex II (atpeninA5, 20 nM) specific inhibitor is used to discriminate complex I driven respiration (yellow) and complex II driven respiration (blue) when mitochondria are incubated with NADH and succinate. Heart (n= 9), Liver (n= 12), error bars represent the mean ± SEM (C, D) complex I driven NAD^+^ synthesis flux assessed in permeabilised mitochondria isolated from heart (C) and liver (D), incubated in presence of cytochrome *c* (62.5 µg/ml), complex I substrate NADH (1.2 mM) and/or complex II substrate succinate (Succ, 10 mM). Heart (n=6), liver (n= 9), error bars represent the mean ± SEM. (E, F) Supramolecular organisation of heart and liver RC. Heart and liver mitochondria are solubilized with a digitonin/protein ratio of 3:1 (g/g) and RC are resolved with 4-16% (E) or 3-12% (F) BN-PAGE, followed by complex I-, complex II- and complex IV-IGA assays, or by western blot using antibody toward ND1, SDHC, UQCRC2 or COX1 subunits. Representative of three different experiments. Heart (n= 3), liver (n= 3). (G, H) RC complexes composition of heart (G) and liver (H). Supramolecular organisation of RC is assessed by 2D-BN/SDS-PAGE and immunoblot analyses are performed with OXPHOS cocktail antibodies toward NDUFB8, MTCO1, UQCR2 and ATP5A subunits. Densitometric profile of complex I (black), complex III (green) and complex IV (orange) detected by immunoblot are represented on top of Coomassie staining. Representative of three different experiments. Heart (n= 3), liver (n= 3). (I) Relative amount of complex I, complex II, complex III and complex IV associated in respirasome (complex I-III-IV) or in supercomplexes containing complex I (complex I_x_-III_y_ or complex I_x_-IV_y_) in heart (red bars) and liver (green bars). Proportion of complexes found associated in respirasome is determined by densitometric quantification of 2D-BN/SDS-PAGE. Heart (n= 3), Liver (n= 3), Error bars represent the mean ± SEM.

The finding that liver and heart RC preferentially oxidize succinate over NADH was in stark contrast to recent hypothesis supporting the fact that the structural association of complex I with complex III and IV should confer kinetics advantage to NADH oxidation over electrons provided by the respirasome free complex II (Enríquez, 2016; Lapuente-Brun et al., 2013). This surprising result prompted us to investigate the supramolecular organization of heart and liver RC. To this end, mitochondria were solubilized using digitonin (at a ratio of 3 g digitonin per g protein) and the supramolecular organization of the RC was then resolved using standard BN-PAGE technique and analysed by western blotting or in-gel activity assay (IGA) (Figure 2E-F and S2E-G). First, BN-PAGE profiles presented in the Figure 2E and 2F clearly confirmed the striking enrichment of heart RC in complex I and IV compared to liver RC, supporting our previous observations (Figure 1E-G). Interestingly, these changes in RC complexes stoichiometry between the two tissues remarkably correlate the abundance and diversity of respirasomes (IxIIIyIVz) (Figure 2E-F). To determine the proportion of complex I and II assembled in supercomplexes, BN-PAGE first dimension was further resolved in a second denaturing dimension. 2D-SDS-PAGE is commonly used to properly quantify the protein levels as they prevent potential epitope accessibility issues encountered with native proteins. Remarkably, 2D electrophoresis analyses were in agreement with previous studies (Greggio et al., 2017; Schägger and Pfeiffer, 2001) showing that more than 90% of heart and liver complex I are found associated in respirasomes whereas no detectable complex II could be found associated in supercomplexes (Figure 2E-I and S2E-G). Furthermore, the supramolecular organization of complex I and II with other RC is not plastic as no significant respirasome reshuffling could be observed when BN-PAGE experiments were performed solubilizing mitochondria actively metabolizing different respiratory substrates (Figure S2H-I). Altogether, our results indicate that despite the absence of stable physical interaction between complex II and other RC complexes, succinate oxidation is prioritized over electrons delivered by complex I.

### Internal oxaloacetate level orchestrates the respective contribution of complex I and II to respiration

To evaluate potential metabolic outcomes of the preferential succinate oxidation on TCA and MAS cycle activities, we applied our new experimental strategy deciphering the origin of electrons fuelling the RC on intact and metabolically active mitochondria. In intact mitochondria, succinate is transported through dicarboxylate carrier before being directly oxidized by complex II. In contrast, NADH cannot be directly imported and will have to be metabolically generated from matrix dehydrogenases like malate, glutamate or pyruvate dehydrogenases. Use of specific inhibitor demonstrated that mitochondrial respiration assessed in presence of pyruvate, glutamate, and malate (PGM) totally relied on complex I activity as this respiration was abolished by rotenone (Figure S2A and C). Similarly, succinate driven respiration was inhibited in presence of AtpeninA5 (Figure S2B and D). Interestingly, uncoupled respiration assessed in presence of succinate and rotenone (Figure 3A-B) were almost identical to succinate driven respiration measured on permeabilized mitochondria (Figure 2A-B). However, when succinate driven uncoupled respiration was assessed in absence of rotenone, respiration values were decreased by 40% in heart and 30% in liver mitochondria (Figure 3A-B and S1C-D). The necessity to add rotenone to assess maximal succinate driven respiration on metabolically active mitochondria has been explained as complex I inhibition could prevent accumulation of oxaloacetate (OAA) a potent competitive inhibitor of complex II (Fink et al., 2018; Gnaiger, 2009; Schollmeyer and Klingenberg, 1961). To characterize the specificity and the metabolic relevance of OAA inhibition, permeabilized mitochondria incubated with succinate or NADH were recorded in presence of increasing concentration of OAA (Figure S3A). These analyses clearly demonstrated that OAA is a powerful and highly specific complex II inhibitor, as no inhibitory side effects could be observed on NADH driven respiration. To elucidate the potential regulatory function of OAA on succinate respiration in intact mitochondria, we developed an enzymatic assay to quantify OAA levels (Figure 3C-D). Our analyses performed on both heart and liver mitochondria confirmed that the stimulation of succinate respiration by rotenone (Figure 3A and B) was associated with a striking lowering of OAA level (Figure 3C-D). As expected, extemporaneous fluorometric measurement of mitochondrial internal NAD(P)H level demonstrated that inhibition of complex I by rotenone massively increased mitochondrial redox (NAD(P)H) state (Figure 3E and S3B). Interestingly, NMR spectroscopy analyses showed that treatment of succinate-metabolizing mitochondria with rotenone, provoked a massive excretion of malate and fumarate (Figure 3F). Altogether, our analyses support that the massive increase in NADH/NAD^+^ ratio secondary to complex I inhibition by rotenone could prevent MDH2 driven OAA accumulation (Figure 3C-D) which consequently increases succinate respiration (Figure 3A-B).

**Figure 3:**
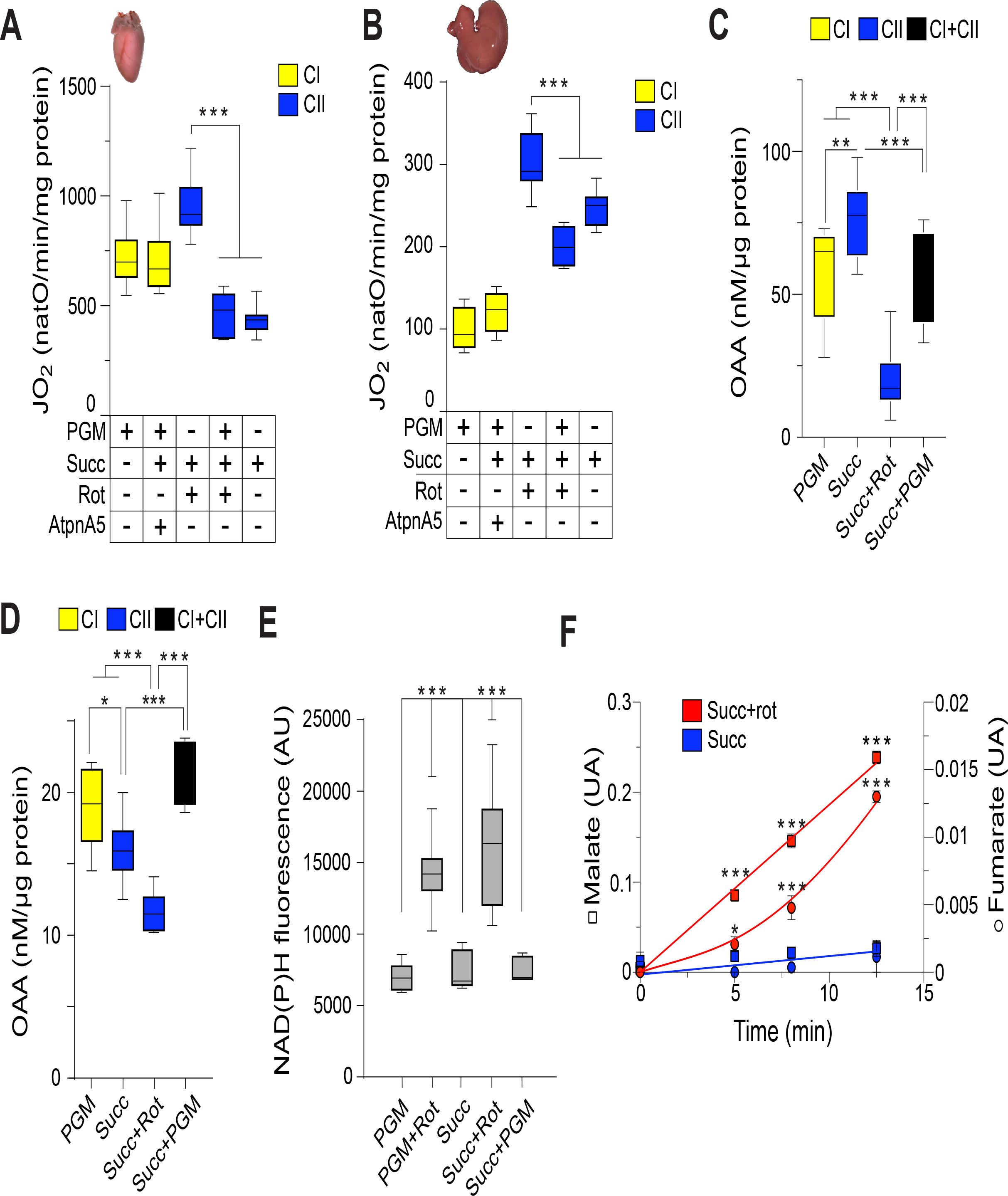
Internal OAA level orchestrates the respective contribution of complex I and II to feed the respiratory chain with electrons. (A, B) Oxygen consumption of uncoupled heart (A) and liver (B) intact mitochondria assessed in presence of complex I substrates (PGM) and/or complex II substrate (Succ). Addition of complex I (rotenone, 30 nM) or complex II (atpeninA5, 20 nM) specific inhibitor is used to discriminate complex I dependent respiration (yellow bars) and complex II dependent respiration (blue bars) when mitochondria are incubated with PGM and Succ. Heart (n= 17), liver (n= 6), error bars represent the mean ± SEM. (C, D) Oxaloacetate levels assessed when uncoupled heart (C) and liver (D) intact mitochondria are fed with complex I substrates (PGM) and/or complex II substrate (Succ). Heart (n= 4), Liver (n= 3), Error bars represent the mean ± SEM. (E) Mitochondrial NAD(P)H fluorescence of uncoupled heart mitochondria incubated with complex I substrates (PGM) ± rotenone (30 nM), or complex II substrate (Succ) ± rotenone (30 nM), or both (Succ+PGM). (n= 5), error bars represent the mean ± SEM. (F) 1H-1D NMR quantification of fumarate (circle) and malate (scare) content are assessed in uncoupled heart mitochondria oxidizing succinate in absence (blue) or in presence of rotenone (30 nM) (red). (n= 3), error bars represent the mean ± SEM.

To determine the respective contribution of complex I and II in fuelling RC with electrons on intact mitochondria, we applied the inhibitor-driven approach. This approach was validated on intact mitochondria as rotenone and atpeninA5 sensitive respirations, assessed under uncoupled conditions, were additive and independent of the sequence of addition of inhibitors (figure S3C-D). Intriguingly, in contrast to the results obtained on permeabilized mitochondria (Figure 2A-B), complex I dependent respiration was preserved in presence of multiple substrates, whereas complex II respiration was similar or even lower than the ‘OAA inhibited’ succinate respiration measured in absence of rotenone (Figure 3A-B). Interestingly, the OAA levels perfectly matched the degree of inhibition of the succinate respiration when all substrates were provided to heart (Figure 3C) and liver (Figure 3D) mitochondria. Altogether, our data identified OAA as a key metabolic regulator orchestrating the respective contribution of complex I and II to RC electron flow. The OAA regulation occurring in intact mitochondria could totally counteract the natural RC priority given to complex II, to favour NADH oxidation.

### MDH2 is a metabolic switch rewiring respiratory chain and TCA fuelling

The importance of MDH2-generated OAA in orchestrating mitochondrial NADH or succinate oxidation, prompted us to investigate the metabolic relevance of this new regulatory mechanism. MDH2 is a mitochondrial metabolic checkpoint located at the crossroad of TCA cycle and MAS (Figure 4A). To decipher the functional interplay between MAS, RC and TCA cycle, we assessed complex II driven respiration mimicking increasing cytosolic NADH reoxidation demand by increasing the extramitochondrial malate level within physiologically relevant concentration range (Figure 4A) (Hautecler et al., 1994; Safer et al., 1971; Siess et al., 1978).

**Figure 4:**
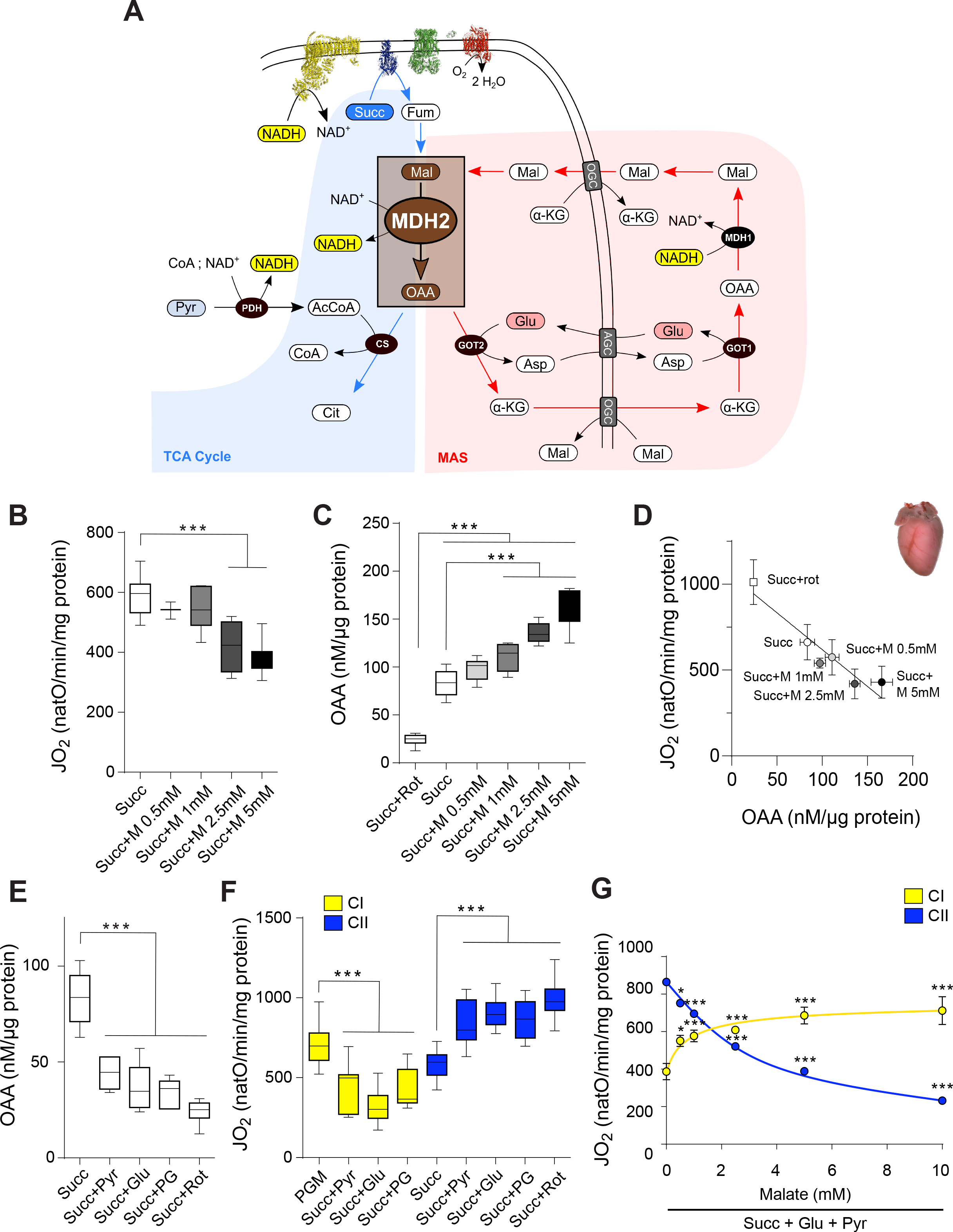
MDH2 rewires TCA cycle and respiratory chain electron flows to favour NADH oxidation. (A) Scheme of respiratory chain, TCA cycle (blue background), and malate-aspartate shuttle (MAS, red background) enlightening the key metabolic role played by MDH2. <u>Abbreviations</u>: succinate (Succ), fumarate (Fum), malate (Mal), oxaloacetate (OAA), glutamate (Glu), aspartate (Asp), α -ketoglutarate (α-KG), acetyl-CoA (AcCoA), citrate (Cit), pyruvate (Pyr), aspartate-glutamate carrier (AGC) 2-oxoglutarate carrier (OGC), malate dehydrogenase (MDH1, MDH2), glutamate-oxaloacetate transaminase (GOT1,GOT2), citrate synthase (CS), pyruvate dehydrogenase (PDH) (B, C) Oxygen consumption (B) and oxaloacetate level (C) assessed with intact heart mitochondria fed with succinate and increased concentration of malate (M) during uncoupled respiration. (n= 4), error bars represent the mean ± SEM. (D) Correlation between mitochondrial oxygen consumption and oxaloacetate levels of intact heart mitochondria isolated fed with succinate and increased concentration of malate (M). Error bars represent the mean ± SEM. (E) Oxaloacetate levels assessed with intact heart mitochondria in presence of different substrates condition metabolizing endogenously produced OAA (Succ+Glu, Succ+Pyr, Succ+PG). (n= 4), error bars represent the mean ± SEM. (F) Oxygen consumption of intact heart mitochondria fed with indicated substrates (Succ+Glu, Succ+Pyr, Succ+PG). Addition of specific inhibitor of complex I (rotenone, 30nM) and complex II (AtpeninA5, 20nM) during uncoupled respiration, determine the complex I driven respiration (yellow bars) or complex II driven respiration (blue bars). (n= 8), error bars represent the mean ± SEM. (G) Correlation between oxygen consumption of intact heart mitochondria fed with succinate, pyruvate, glutamate and increasing concentration of malate. Complex I driven (yellow circle) or complex II driven (blue circle) respiration are determined through sequential addition of specific complex I (rotenone, 30nM) and complex II (AtpeninA5, 20nM) inhibitors. (n= 4), error bars represent the mean ± SEM.

This experiment demonstrated that complex II driven respiration decreased proportionally in response to increased malate concentration (Figure 4B-D and S4A-C). As control, we showed that malate *per se* did not impact succinate respiration in permeabilized mitochondria (Figure S4D). This observation further explained why malate inhibition only occurred in metabolically active mitochondria where OAA can be generated and accumulated. In line with this observation, the decrease in complex II driven respiration associated with increase malate concentration was perfectly correlated with an increase in mitochondrial OAA level (Figure 4D and S4C). Altogether, our data show that more than 60% of complex II driven respiration can be downregulated in response to increased extramitochondrial malate concentration through a MDH2-mediated metabolic regulation in both heart and liver.

To confirm that OAA could act as mitochondrial metabolic sensor adjusting RC fuelling to mitochondrial metabolic activity, we investigated how OAA consuming pathways impact succinate and NADH dependent respirations. To this end, mitochondrial RC activity was assessed in presence of succinate (Succ) and glutamate (Glu) or/and pyruvate (Pyr) to metabolically flush OAA through MAS (GOT2 dependent pathway) or through the TCA cycle (pyruvate dehydrogenase and citrate synthase dependent pathway) (Figure 4A). Our analyses demonstrated that both pyruvate and glutamate were efficient in consuming internal OAA as both substrates added alone or in combination decreased OAA to similar residual level as the one measured in presence of succinate and rotenone (Figure 4E and S4E). The metabolic flush of OAA did not only increase succinate driven respiration by releasing complex II inhibition, but it also fuelled complex I with NADH (Figure 4F and S4F). Eventually, we decided to determine if MDH2 could metabolically rewire RC electrons entry from succinate toward NADH, when MAS driven external NADH oxidation demand is increased (increasing malate concentration within physiological range). Our results clearly showed that Malate-OAA dependent respiration rewiring from complex II to complex I occurs, even when OAA produced by the MDH2 was metabolically flushed through MAS and TCA cycle (Figure 4G). Remarkably, imported malate oxidized through MDH2 not only downregulates succinate driven respiration, but it also increases MDH2 driven NADH production and complex I driven NADH oxidation capacity.

## DISCUSSION

Our present work indicates that the differences between heart and liver mitochondrial bioenergetics activities mainly correlate with change in abundancy and stoichiometry of OXPHOS complexes. For instance, differences in complex I and II levels (Figure 1H) are in line with the different NADH and succinate driven respiration capacities of heart and liver mitochondria (Figure 1A-D). In contrast, the proportion of complex I and II assembled into respirasome is not changed between heart and liver RC (Figure 2E-I and S2E-I). In line with previous works performed in different tissues or model organisms, the supramolecular organisation of mouse heart and liver RC demonstrate that about 90% of complex I is associated in supercomplexes whereas complex II is never found associated with other RC complexes (Greggio et al., 2017; Schägger and Pfeiffer, 2001). This striking difference in the supramolecular organisation of the CoQ-oxidoreductases led to the assumption that the physical association of complex I with other RC complexes could reduce the diffusion distance of CoQ and Cyt c between organized complexes, promoting NADH oxidation (Balsa et al., 2019; Bianchi et al., 2004; Lapuente-Brun et al., 2013). To decipher the functional relevance of the supramolecular organization of the complex I and II on their ability to fuel the RC, we developed a new experimental strategy using permeabilized mitochondria to (i) provide non-limited amount of NADH and succinate and follow their oxidation rate, (ii) remove thermodynamic constraints linked to the protonmotive force on complex I and on the RC activity, (iii) prevent the interfering inhibition of complex II by OAA. Surprisingly, the inhibitor driven approach as well as multiple flux (JNAD^+^ and JO2) analyses consistently demonstrates that when permeabilized mitochondria are incubated with both NADH and succinate, complex II driven respiration is preserved whereas complex I driven respiration is strongly and significantly decreased in heart and liver mitochondria (Figure 2A-D). Interestingly, the fact that NADH driven respiration is drastically reduced in the presence of succinate clearly demonstrates that the respirasome is not functionally disconnected from the respirasome free complex II (Figure 2C-D). Our observations indicates that respirasomes structuration does not lead to strict, physical and functional, partitioning between RC complexes and electrons carriers (CoQ and Cyt c) and thus support previous work (Bianchi et al., 2004; Blaza et al., 2014; Fedor and Hirst, 2018). The discovery of a mechanism enabling the RC to orchestrate the activities of coenzymes Q oxidoreductases constitutes a major conceptual breakthrough in the field of mitochondrial energy metabolism. This regulatory mechanism called ‘electron competition’ or ‘CoQ-contest’ originally discovered in *S.cerevisiae* allow the yeast RC to prioritize oxidation of cytosolic NADH (Bunoust et al., 2005; Mourier et al., 2010; Påhlman et al., 2001). The fact that, in contrast to succinate oxidation, the NADH driven respiration is found drastically affected when heart and liver RC are incubated with both NADH and succinate clearly demonstrate that the ‘CoQ contest’ mechanisms is not restricted to the yeast model but also orchestrates mammalian heart and liver RC electron flow (Figure 2A-D). Moreover, in line with observations made with yeast, the priority given to certain CoQ-oxidoreductase is (i) independent of the RC supramolecular organisation (Rigoulet et al., 2010) and (ii) independent of the stoichiometry between CoQ-oxidoreductases involved (Figure 1F-H and Figure 2E-F).

The paramount importance of the maintenance of the NADH/NAD^+^ redox balance by the mitochondria for cell metabolism and cell functions was consistently observed in various eukaryotes models expressing different fermentative pathways and presenting different NADH CoQ-oxidoreductase systems fuelling the RC (Carneiro et al., 2004; Foriel et al., 2019; Grad and Lemire, 2004; Luttik et al., 1998). In humans complex I or redox shuttles loss of functions cause numerous pathologies and the resulting NADH redox imbalance often lead to life-threatening lactic acidosis crises (Ait-El-Mkadem et al., 2017; Hayes et al., 1987; Munnich and Rustin, 2001; van Karnebeek et al., 2019). Recent studies have shown that the control of the cellular redox balance by the mitochondria is essential for the maintenance of cellular proliferative capacities (Birsoy et al., 2015; Sullivan et al., 2015). Until recently, mitochondria were described as being impermeable to NAD^+^/NADH (Purvis and Lowenstein, 1961). However, this dogma has been recently challenged by the demonstration that NAD(H) could be imported in mammalian mitochondria (Davila et al., 2018) and by the identification of SL25A51 as a mammalian mitochondrial NAD(H) transporter (Luongo et al., 2020) compensating the absence of intramitochondrial NAD(H) synthesis. However, the rate of NADH import through this transporter is extremely limited and therefore is not directly involved in connecting cytosolic and mitochondrial NADH/NAD^+^ redox homeostasis. Instead, connection between the cytosolic and mitochondrial redox homeostasis is ensured by redox shuttles such as the MAS (LaNoue et al., 1974, 1973).

The impressive enzymatic equipment metabolizing cytosolic and mitochondrial OAA demonstrates its metabolic importance and positioned this metabolite at the crossroad between glycolysis (Pyruvate Carboxylase 1 and 2), gluconeogenesis (PEPCK1, PEPCK2), TCA cycle (MDH2, CS) and Malate aspartate Shuttle (MDH1, MDH2, GOT1, GOT2). Bioenergetic analyses of heart and liver mitochondria, early on characterized through their high MAS metabolic capacities (LaNoue et al., 1973; Safer B, 1975), support recent work from the Sivitz group claiming that regulation of complex II activity by OAA is of great metabolic interest (Fink et al., 2019, 2018). Our analyses extend and refine the mitochondrial metabolic processes controlling internal OAA levels and therefore regulating the complex II activity. Importantly, our data demonstrate that MDH2 acts as a metabolic conductor synergistically orchestrating RC and TCA activity to maximise mitochondrial NADH oxidation capacity. First, our work confirms that production of OAA by MDH2 relies on mitochondrial redox (NADH/NAD+) maintenance by complex I and can be abolished when complex I is inhibited by rotenone (Figure 3C-F and S3B). Secondly, when malate is not present and imported, the complex II inhibition caused by MDH2 driven OAA production can be completely reversed when OAA is metabolized by PDH and CS (succ+pyr) or GOT2 (succ+glut) (Figure 4E-F). Surprisingly, we show that extra-mitochondrial malate, added at physiologically relevant concentrations range, can increase internal OAA, leading to a dominant inhibition of complex II, rewiring RC electrons fuelling as well as the TCA anaplerosis. This dose dependent inhibition of the complex II driven respiration by extramitochondrial malate is associated with an increase in complex I driven respiration (Figure 4G). Beyond complex II inhibition, MDH2 activity is crucial to sustain complex I activity through direct NADH production and by producing OAA indispensable for mitochondrial metabolism of pyruvate and glutamate (Figure S1A-B).

Moreover, the immediate and reversible inhibition of complex II by OAA indicate that mitochondrial OAA levels could act as a metabolic sensor adjusting electrons fuelling the RC to fit cell metabolic needs. This metabolic regulation synergistically increases RC’s NADH oxidative capacity and rewires MDH2 driven TCA anaplerosis, preventing malate production from succinate to favour oxidation of imported malate supporting MAS activity (Figure 4G). This discovery prompt us to hypothesise that MAS does not only passively balance cytosolic and mitochondrial NADH but instead, through MDH2 oxidation of imported malate, MAS actively rewires RC’s fuelling, and TCA anaplerosis to boost complex I activity and fuelling.

## EXPERIMENTAL PROCEDURES

### Biological materials

All mice used in this study were on an inbred C57Bl/6N background. Mice were maintained on a standard mouse chow diet and sacrificed, between 20 and 40 weeks of age, by cervical dislocation in strict accordance with the recommendations and guidelines of the Federation of the European Laboratory Animal Science Association, and obtain with an authorization from the French ministry of Agriculture (APAFIS#12648-2017112083056692v7).

### Heart and liver mitochondria Isolation

Isolation of heart and liver mitochondria was performed by differential centrifugation as previously described (Brandt et al., 2017; Mourier et al., 2015). Briefly, right after the sacrifice, tissues were collected, minced and cleaned with mitochondria isolation buffer (MIB; 310 mM sucrose, 20 mM Tris-Base, 1 mM EGTA, pH 7.2) for the heart and MIB+BSA (MIB and 0.25mg/ml of delipidated Bovine serum albumin (BSA)) for the liver. The pieces of heart were collected and homogenized with few strokes of a Potter S homogenizer (Sartorius) using a loose Teflon pestle, in an ice cold MIB supplemented with trypsine (0.5 g/l). Liver tissues homogenisation was performed in a Potter S homogenizer (Sartorius) using a Teflon pestle, and in ice cold MIB+BSA. Mitochondria were isolated by differential centrifugation, a low speed one (1,000g, 10 min, 4C) in a swing out rotor and another one at high speed (3,500g, 10 min, 4°C for heart mitochondria and 10,000g, 10 min, 4C for liver mitochondria) using a fixed angle rotor. The pellet of crude mitochondria obtained is suspended in MIB+BSA. Protein concentration was measured using Bio-Rad DC Protein Assay kit according to provider’s instructions.

### High-resolution oxygen consumption measurement

Oxygen consumption of intact crude mitochondria was measured at 37°C, using respectively 50µg and 100µg of heart and liver mitochondria diluted in 2.1ml of respiratory buffer (RB : 120 mM sucrose, 50 mM KCl, 20 mM Tris-HCl, 4 mM KH2PO4, 2 mM MgCl2, 1 mM EGTA, 0.25 mg/ml delipidated BSA; pH 7.2) in an Oxygraph-2k (OROBOROS INSTRUMENTS, Innsbruck, Austria) (Mourier et al., 2014). Oxygen consumption was measured by using different association of substrates: succinic acid (5 mM; pH7.2), sodium pyruvate (10 mM; pH7.2), L-Glutamate acid (10 mM; pH7.2), L-Malic acid (concentrations are indicated in the figure legends; pH7.2). Oxygen consumption rate was measured under 3 different conditions: in the phosphorylating state with addition of ADP (5 mM; pH7.2), in the non-phosphorylating state with addition of oligomycin (50 ng/ml), and in the uncoupled state by successive addition of carbonyl cyanide m-chlorophenyl hydrazone (CCCP) up to 0.5 µM to reach the maximal respiration of the RC.

Activity of the RC was also recorded using permeabilized crude mitochondria obtain after freeze and thaw cycle. Oxygen consumption by the RC was measured as previously described (Mourier et al., 2014), using respectively 50 and 100µg of heart and liver mitochondria diluted in 2.1ml of potassium phosphate buffer (50 mM, pH 7.2) and incubated with saturating concentrations of cytochrome *c* (62.5 µg/ml), NADH (0.625 mM) and succinate (10 mM). No oxygen consumption could be detected in absence of cytochrome *c*, and no oxygen consumption link to complex I activity was recorded when permeabilized mitochondria are incubated with pyruvate, glutamate and malate, confirming the loss of integrity of mitochondrial internal compartment.

The determination of oxygen consumption link to electron coming from complex I and complex II was determined after addition of rotenone (Rot, 30 nM) for complex I and/or atpenin A5 (AtpnA5, 20nM) for complex II. As a control, Antimycin A (100 nM) was added at the end of each experiment to control the proportion of oxygen consumption which is not linked to RC.

### Mitochondrial cytochromes quantification

The different cytochromes of the mitochondrial respiratory chain were measured by dual-wavelength spectrophotometry by comparing the spectra of fully oxidized *vs*. fully reduced cytochromes. In each of the two cuvettes, 2 mg of heart or 16 mg of liver mitochondrial protein were suspended in 1 ml of phosphate buffer (50 mM, pH 7.2), and 0.05% (vol/vol) Triton X-100 were added. In the “oxidized” cuvette, 10 µl of ferricyanide (0.5 M) were added while a few grains of sodium hydrosulfite were added in the “reduced” cuvette. The components were mixed, and the absorbance spectrums of both cuvettes were recorded. A typical difference between reduced minus oxidized spectra was obtained. Wavelength pairs and absorption coefficient used were: cytochrome *c* + *c1* (550–540 nm) ε = 18 mM^-1^/cm, cytochrome *b* (563– 575 nm) ε = 18 mM^-1^/cm and cytochrome *a* + *a3* (605–630 nm) ε = 24 mM^-1^/cm.

### Western-blot analyses

Proteins from crude heart and liver mitochondria were resolved using denaturing (SDS PAGE or native electrophoreses (BN-PAGE and Second dimensional SDS PAGE)).

For SDS PAGE, 50 µg of heart and liver crude mitochondria proteins were solubilised with RIPA buffer (NaCl (150 mM), tris-base (25 mM), NP40 (1% w/v), SDS (1% w/v), deoxycholate (0.25% w/v) and EGTA (1 mM), pH 8). Solubilised mitochondria were mixed with Laemmli buffer (Laemmli, 1970), separated by Bis-Tris 15% acrylamide gel, and then transferred on nitrocellulose Amersham Protran Premium membrane (Amersham). Immunodetection was performed by fluorescence using a Typhoon FLA 9500 (GE Healthcare Life Sciences). To perform immunodetection, membranes were blocked with 3% (w/v) milk diluted in Tris Buffer Saline (TBS; pH 8), then primary antibody (Total OXPHOS Rodent WB Antibody Cocktail; Abcam) was added for incubation, and finally an ECL Plex™ G-A-R IgG, Cy^®^5 (GE Healthcare) anti-mouse secondary antibody was used. The FIJI software was used to determine and analyse densitometric profiles.

### BN-PAGE analyses

For BN-PAGE, 50 µg of crude heart mitochondria and 150 µg of crude liver mitochondria proteins were incubated with digitonin extraction buffer (HEPES (30 mM), potassium Acetate (150 mM), Glycerol (12%), 6-Aminocaproic acid (2 mM), EDTA (2 mM), high-purity digitonin (3 g/g), pH 7.2), and vortexed 1h at 4°C to solubilize membranes. After incubation, mitochondria were centrifugated at 30,000 g for 20 minutes, supernatants were collected and mixed with loading dye (0.0125% (w/v), Coomassie brilliant blue G-250). Native complexes were resolved using Bis-Tris Invitrogen™ Novex™ NativePAGE™ 3-12% or 4-16% acrylamide gradient. To reach the optimal separation and resolution of OXPHOS supramolecular organisations, digitonine solubilised proteins were loaded on 3-12% gradient gel and were migrated at 10 mA for 12 h at 4°C (complex II are not retained in the gel after overnight run). To analyse the complex II assembly, 4-16% gradient gels were migrated 6 h in order to keep the complex II in gel and for optimal separation of RC supercomplexes. RC complexes were detected using in-gel activity (IGA) assays as described previously (Wittig et al., n.d.). Briefly, native gels were washed and incubated in a potassium phosphate buffer (50 mM, pH 7.2) containing iodonitrotetrazolium (1 mg/ml) supplemented with NADH (400 µM, pH7.2) for complex I or Succinate (50 mM, pH7.2) and Phenazine Methosulfate (20mM) for complex II. For complex IV in-gel activity assays, native gels were incubated in RB containing Diaminobenzidine (1 mg/ml) and cytochrome *c* (0.5 mg/ml). Coomassie staining was performed with PageBlue protein staining solution. Colorimetric bands were recorded with Optical Densitometry of an Amersham ImageQuant 800.

Proteins were also immunodetected, gels were incubated in Towbin buffer (Tris (0.3% w/v), Glycine (1.44% w/v), Ethanol (10% v/v)) supplemented with SDS (0.2% v/v) and β-mercaptoethanol (0.2% v/v) for 30 minutes at RT, to denature proteins. Then gels were transferred on polyvinylidene difluoride (PVDF) in Towbin transfer buffer for 2h at 0.2 A. Immunodetection was performed as described previously, peroxidase-conjugated anti-mouse and anti-rabbit IgGs and Amersham CyDye 800 goat anti-rabbit and anti-mouse were used for immunodetection by chemiluminescence and fluorescence. Immunoreactive bands were recorded with an Amersham ImageQuant 800.

BN-PAGE gels after first dimension electrophoresis, were ran in a denaturing second dimension electrophoresis (2D-SDS PAGE). After collecting the first-dimension band and before the second dimension, first dimension bands were incubated in an Invitrogen™ Bolt™ MOPS SDS Running Buffer (Fisher Scientific) supplemented with β-mercaptoethanol (1% v/v) for 30min at RT. Acrylamide gradient Bolt™ 4-12% Bis-Tris Plus Gels (Thermo Scientific) were used for 2D SDS-PAGE, the gels were transferred and immunodetection was performed as previously described for classical SDS-PAGE. Immunoreactive bands were recorded by fluorescence and chemiluminescence with an Amersham ImageQuant 800. The FIJI software was used to quantify the perform densitometry analyses of images obtained.

### Oxaloacetate quantification

Oxaloacetate level was measured using enzymatic assay based on citrate synthase activity. Intact heart (200µg) and liver (1000µg) mitochondria were incubated like for oxygen consumption measurement in an Oxygraph-2k at 37°C. Briefly, 150 µl of mitochondria incubated in 2.1 ml RB consuming oxygen under uncoupled condition were collected and quenched with 50µl ice cold acidic perchloric acid (PCA 7% without EDTA). Samples were then centrifuged 10 minutes at 30,000g and supernatants were quickly neutralized with KOMO (KOH (2 M); MOPS (0.5 M)). OAA content was then quantified using citrate synthase buffer (Acetyl-CoA (200 µM), 5,5’-dithiobis-(2-nitrobenzoic acid) (200 µM), Citrate Synthase from pig heart (4 U/ml) in phosphate buffer (50 mM, pH 7.2). Absorbance was recorded at 412 nm in UV Greiner bio-one 96-well plate by a CLARIOstar plate reader (BMG Labtech). Standard curve allowing OAA quantification was determined using fresh OAA solution.

### NADH oxidation flux measurement (JNAD^+^)

NAD^+^ production rate was measured using enzymatic assay using G6PDH from *leuconostoc mesenteroides.* permeabilized mitochondria from heart (50 µg) and liver (100 µg) were incubated in 2.1 ml RB in an Oxygraph-2k at 37°C. 150 µl of mitochondria incubated in 2.1 ml RB buffer consuming oxygen under uncoupled condition were collected and quenched with 50µl ice cold acidic perchloric acid (PCA 7% without EDTA). Samples were then centrifuged 10 minutes at 30,000g and supernatants were quickly neutralized with KOMO (KOH (2 M); MOPS (0.5 M)). Neutralized samples were then subjected to enzymatic assay quantifying NAD^+^ content as described previously (Bunoust et al., 2005).

### Fumarate and Malate metabolic flux measurement

Metabolic fluxes were measured by ^1^H-1D NMR. Intact Heart mitochondria were incubated in the same conditions than oxygen consumption measurements in an Oxygraph-2k at 37°C. During steady state respiration assessed under uncoupled conditions, aliquots collected at 0, 5, 7.5, and 12.5 minutes of incubation were snap frozen in liquid nitrogen to quench biochemical reactions. Metabolites were extracted with a mix of acetonitrile (ACN) and methanol (MeOH) to a final ACN/MeOH/Water proportion of 2:2:1. They were then vortexed, incubated at -20C for 20 min, and evaporated with a ThermoFisher SpeedVac SC250EXP concentrator. Samples were then dissolved into 200µL D2O solution containing 2 mM Trimethyl-sillyl-propionic acid d4 (TSPd4) used as NMR reference standard, and centrifuged to remove membrane debris. A final 180 µL volume of the resulting supernatants were transferred into 3 mm NMR tubes. Samples were analyzed by 1H-1D NMR on a Bruker Ascend 800 MHz NMR spectrometer equipped with a 5 mm QPCI cryoprobe. A quantitative zgpr30 sequence was used with 64k acquisition points during 128 scans. The total repetition time between scans was set to 7 seconds. Malate and fumarate concentrations were quantified from their respective signals in the 1H-1D NMR spectra using TSPd4 as internal standard. All measurements were performed in triplicates.

### Statistical analyses

Data are presented as mean ± SEM unless otherwise indicated in figure legends. Sample number (n) indicates the number of independent biological samples (individual mice) for each experiment. Sample numbers and experimental repeats are indicated in the figures. Data were analyzed with the GraphPad Prism software using unpaired Student’s t-test, one-way ANOVA using Turkey’s multiple comparison test, or two-way ANOVA using Bonferroni multiple comparison test between group comparison, as appropriate. A 0.05 p-value was considered statistically significant.

## ACKNOWLEDGEMENTS

We gratefully acknowledge for the support, encouragement and useful discussions provided by Dr. Manuel Rojo. We would like to thank Prof. Michel Rigoulet and Prof. Nils-Göran Larsson, Dr. Dusanka Milenkovic for their precious feedback on the manuscript. We also thank Benoit Rousseau and Julien Izotte for the work at animal house (the service commun des animaleries, Animalerie A2 – University of Bordeaux). This work was supported by the AFM-Telethon Trampoline (AFM-19613) and ANR (ANR-16-CE14-0013). The funders had no role in study design, data collection and analysis, decision to publish, or preparation of the manuscript. MetaboHub-MetaToul (Metabolomics & Fluxomics facilities, Toulouse, France, http://www.metatoul.fr) is supported by the ANR grant MetaboHUB-ANR-11-INBS-0010. JCP is grateful to INSERM for funding a temporary full-time researcher position.

---

**Figure 1S:**
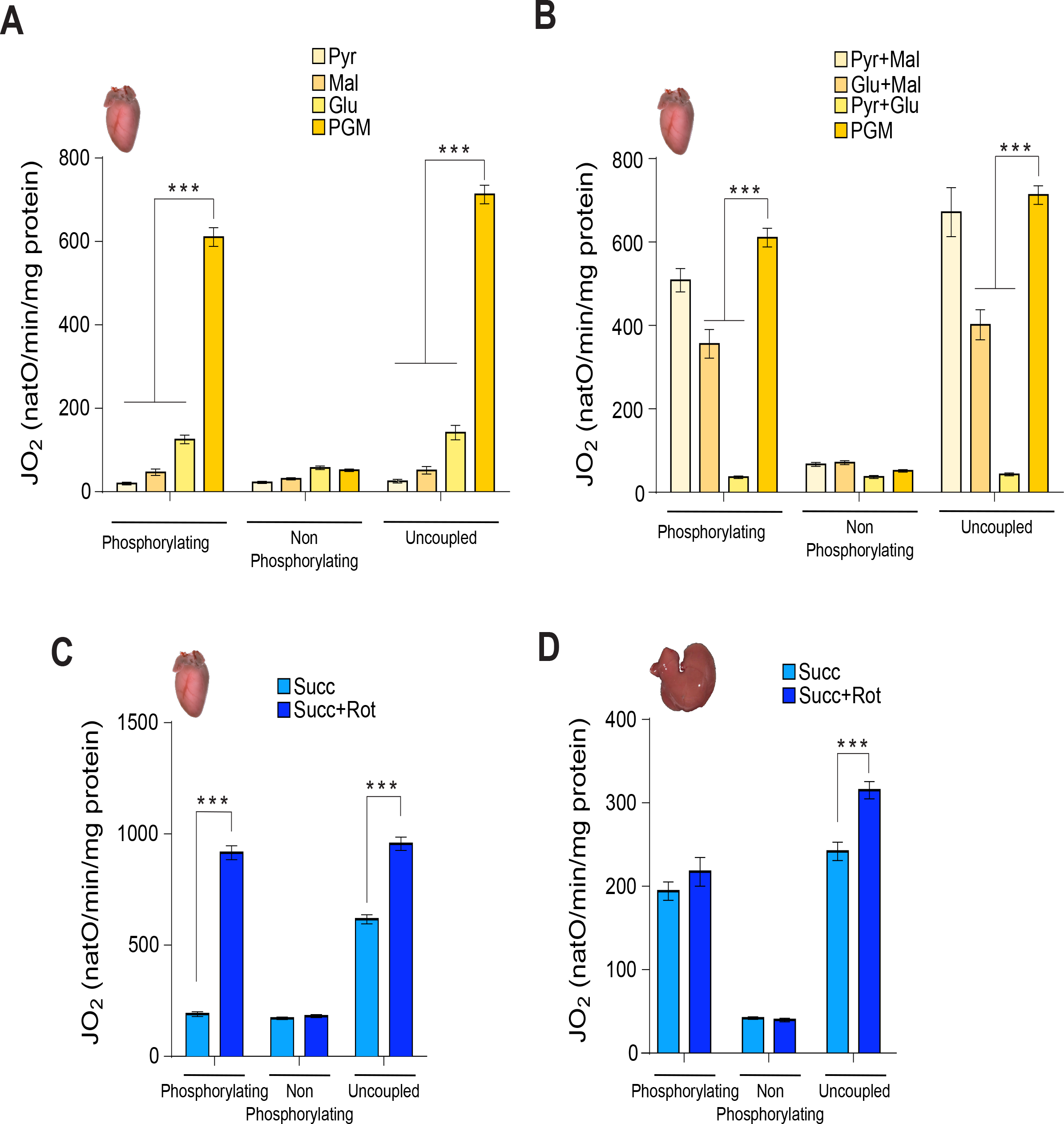
Complex I and complex II driven respiration depends strongly on substrate combination. (A, B) Oxygen consumption of intact mitochondria isolated from heart were assessed in presence of unique (A) or mixed combination of substrates (B) which mitochondrial metabolism result in delivering NADH to complex I (pyruvate 10 mM, glutamate 5 mM, malate 5 mM). Respiration are assessed under phosphorylating conditions (ADP+Pi), non-phosphorylating conditions (oligomycin), and uncoupled (CCCP titration). (n= 4) Error bars represent the mean ± SEM (C, D) Oxygen consumption of intact mitochondria isolated from heart (C) and liver (D) are assessed in presence of succinate ± rotenone (30 nM) under phosphorylating conditions (ADP+Pi), non-phosphorylating conditions (oligomycin), and uncoupled (CCCP titration). Heart (n= 28), Liver (n= 11), Error bars represent the mean ± SEM

**Figure 2S:**
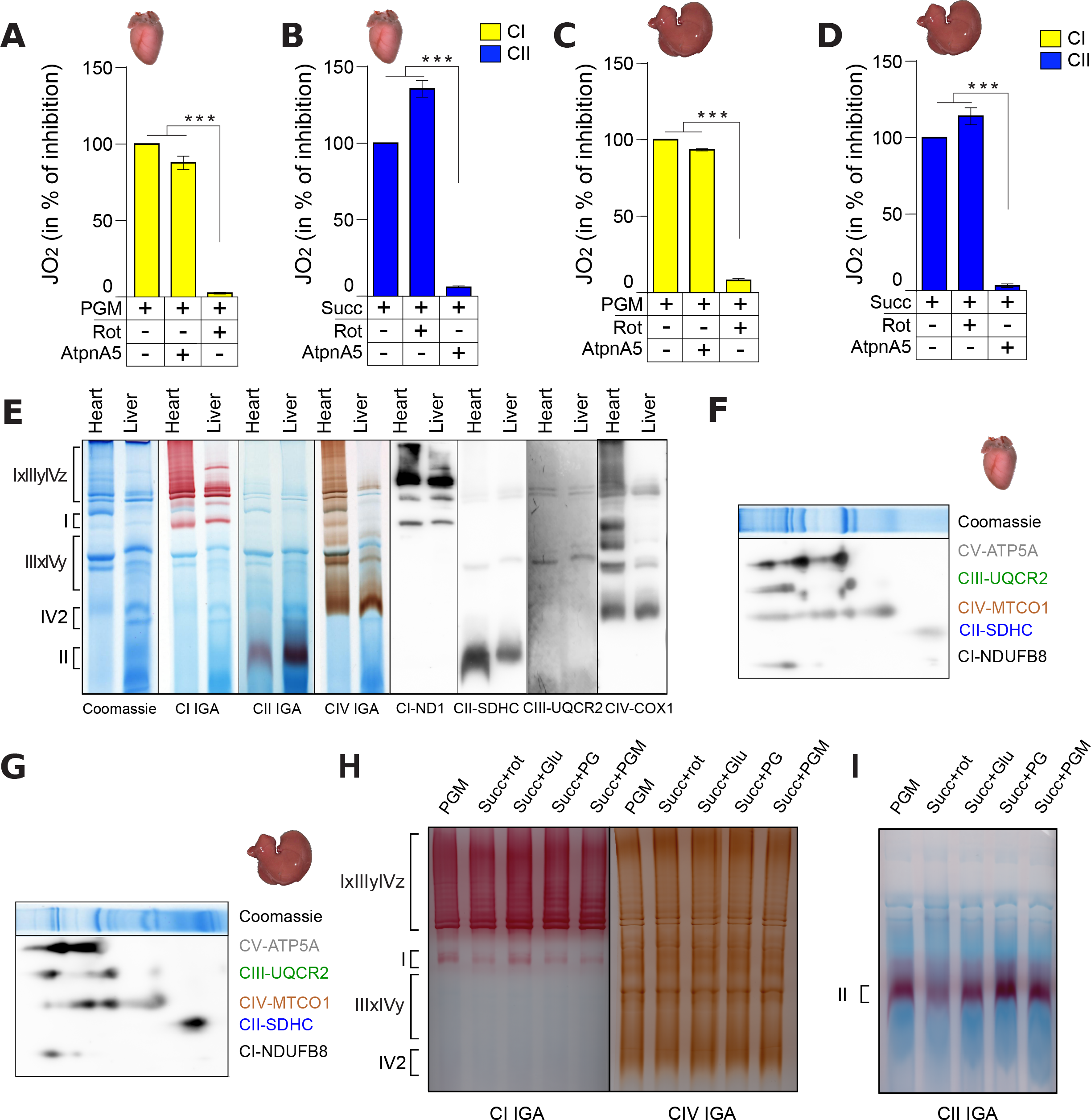
Heart and liver mitochondrial respiratory chain’s enzymes stoichiometry, supramolecular organization and preferential electron fuelling. (A-D) Oxygen consumption of uncoupled heart (A, B) and liver (C, D) intact mitochondria in presence of mitochondrial complex I substrates (PGM, yellow bars) (A, C), or complex II substrate (Succ, blue bars) (B,D). Complex I (A, C) or complex II (B, D) driven respiration are recorded ± rotenone (rot, complex I inhibitor, 30 nM), or Atpenin A5 (AtpnA5, complex II inhibitor, 20 nM). Heart (n= 5), liver (n= 5), Error bars represent the mean ± SEM. (E) Supramolecular organisation of heart and liver respiratory chain. Heart and liver mitochondria are solubilized with a digitonin/protein ratio of 3:1 (g/g) and high MW assemblies are resolved using 4-16% BN-PAGE, followed by complex I-, complex II- and complex IV- IGA assays, or by western blot analyses using antibody against ND1, SDHC, UQCRC2 or COX1 subunits. Representative of three different experiments. Heart (n= 3), liver (n= 3). (F, G) RC complexes composition of heart (F) and liver (G) SC assessed by 2D-BN/SDS-PAGE and immunoblot analyses are performed with OXPHOS cocktail antibodies against NDUFB8, MTCO1, UQCR2 and ATP5A subunits. Heart (n= 3), liver (n= 3). (H, I) Supramolecular organisation of heart respiratory chain incubated with different respiratory substrate. Heart and liver mitochondria incubated with PGM, succinate ± rotenone (Succ, Succ+rot), succinate glutamate (Succ+Glu) or succinate pyruvate and glutamate (Succ+PG) are solubilized with a digitonin/protein ratio of 3:1 (g/g) and RC are resolved with 3-12% (H) or 4-16% (I) BN-PAGE, followed by complex I-, complex II- and complex IV-IGA assays. Representative of three different experiments. Heart (n= 3).

**Figure 3S:**
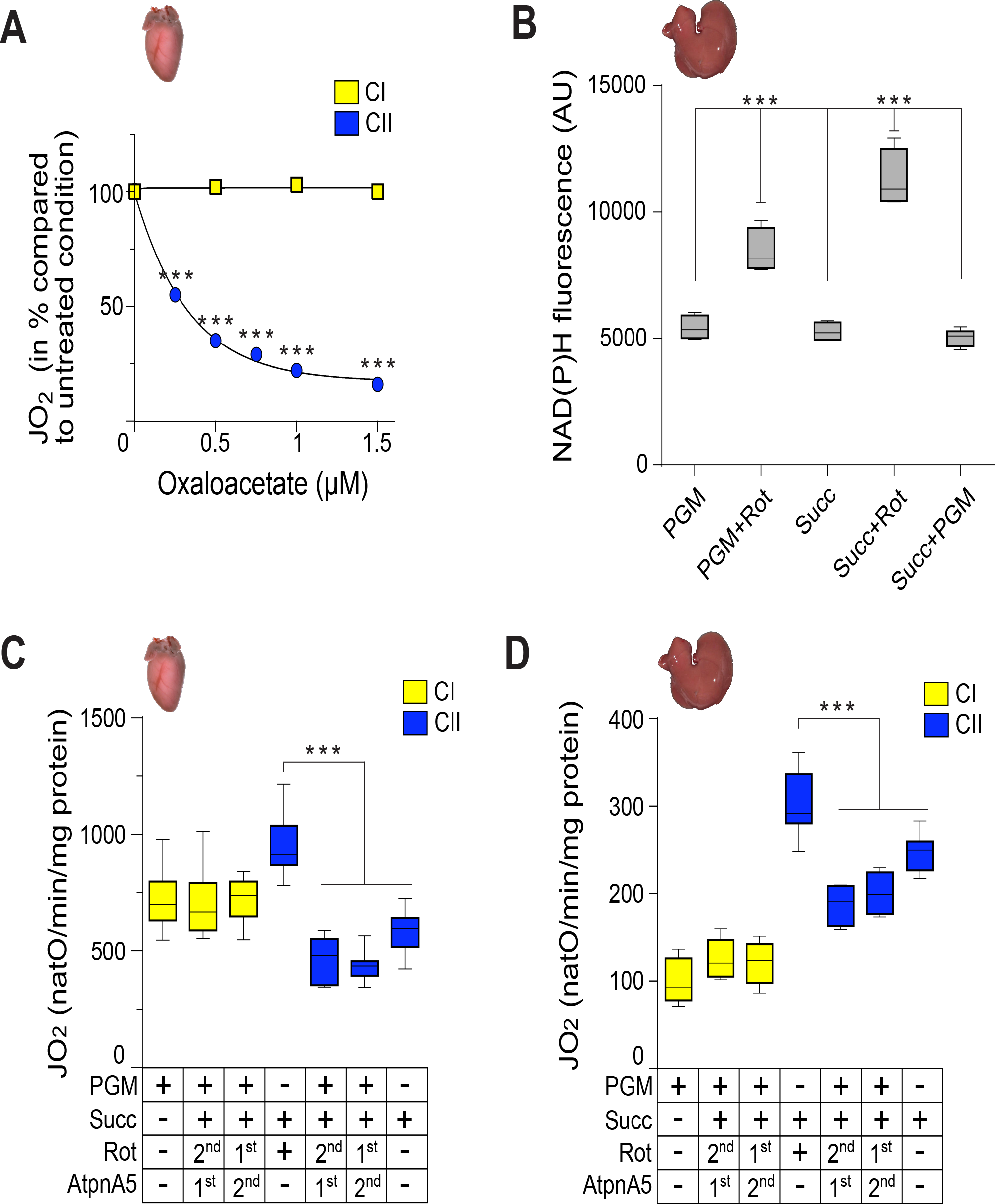
Internal OAA level orchestrates the respective contribution of complex I and II to feed the respiratory chain with electrons. (A) Oxygen consumption of permeabilised heart mitochondria in presence of cytochrome *c* (62.5 µg/ml), and increasing concentration of oxaloacetate are added when mitochondria are incubated with NADH (yellow bars) or with succinate (blue bars). (n=3), error bars represent the mean ± SEM. (B) NAD(P)H fluorescence of uncoupled liver mitochondria incubated with complex I substrates (PGM) ± rotenone (30 nM), or complex II substrate (Succ) ± rotenone (30 nM), or both (Succ+PGM). Liver (n= 3), error bars represent the mean ± SEM. (C,D) Oxygen consumption of uncoupled heart (C) and liver (D) intact mitochondria fed with complex I substrates (PGM) and/or complex II substrate (Succ). Addition of complex I (rotenone, 30 nM) or complex II (atpeninA5, 20 nM) specific inhibitor are used to discriminate complex I dependent respiration (yellow bars) and complex II dependent respiration (blue bars) when mitochondria are incubated with both PGM and succinate. Complex I and complex II inhibitor were added sequentially (sequence addition are indicated in the table below the graph: 1^st^ for firstly added and 2^nd^ for the one added after). Heart (n= 11), liver (n= 4), error bars represent the mean ± SEM.

**Figure 4S:**
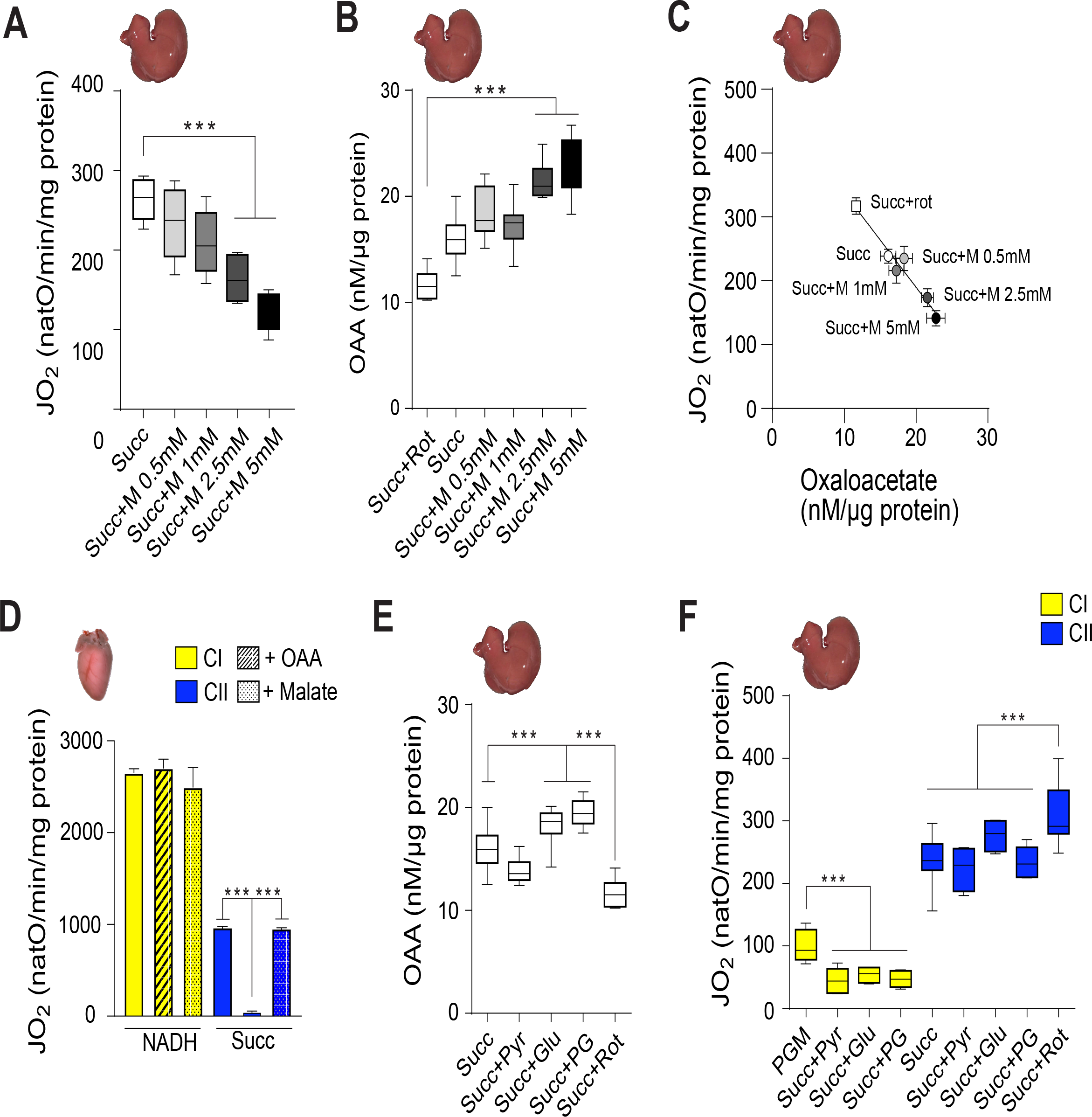
MDH2 rewires TCA cycle and respiratory chain electrons flow to favour NADH oxidation. (A, B) Oxygen consumption (A) oxaloacetate level (C) assessed with intact liver mitochondria fed with succinate and increased concentration of malate (M) during uncoupled respiration. (n= 3), error bars represent the mean ± SEM. (C) Correlation between mitochondrial oxygen consumption and oxaloacetate levels of intact liver mitochondria isolated fed with succinate and increasing concentration of malate (M). Error bars represent the mean ± SEM. (D) Oxygen consumption of permeabilised heart mitochondria fed with complex I (NADH, yellow bars) or complex II (Succ, blue bars) substrates ± addition of oxaloacetate (hatched bars) or malate (dotted bars). (n= 4), error bars represent the mean ± SEM. (E) Oxygen consumption of uncoupled heart mitochondria fed with succinate (10mM) and malate (5 mM). Addition of complex I (rotenone, 30 nM) or complex II (atpeninA5, 20 nM) specific inhibitor are used to quantify the complex I driven respiration (yellow bars) and complex II driven respiration (blue bars). Complex I and complex II inhibitor were added sequentially (sequence addition are mentioned in the table below the graph). (n= 5), error bars represent the mean ± SEM. (F) Oxaloacetate levels assessed with intact liver mitochondria in presence of different substrates condition metabolizing endogenously produced OAA (Succ+Glu, Succ+Pyr, Succ+PG). (n= 3), error bars represent the mean ± SEM. (G) Oxygen consumption of intact liver mitochondria fed with indicated substrates conditions (Succ+Pyr, Succ+Glu or Succ+Pyr+Glu). Addition of specific inhibitor of complex I (rotenone) and complex II (AtpeninA5) during uncoupled respiration, determine the complex I driven respiration (yellow bars) or complex II driven respiration (blue bars). (n= 5), error bars represent the mean ± SEM.

